# Nuclear envelope-dependent heterochromatin positioning gates the timing of nucleolar assembly in the early embryo

**DOI:** 10.64898/2026.07.20.739428

**Authors:** Xiangyi Ding, Guanzhi Wang, Daniel Elnatan, Matthew Laurence, Sara Zdanovskis, Megan Couture, Stephanie C. Weber, Daniel A. Starr, G.W. Gant Luxton

## Abstract

The nucleolus is the largest nuclear condensate, yet how its assembly is developmentally timed remains poorly understood. In the *Caenorhabditis elegans* embryo, nucleoli normally appear at the 6- to 8-cell stage. Here we show that the LINC complex and nuclear lamina restrain premature nucleolar assembly through heterochromatin organization. Depletion of the embryonic LINC complex proteins SUN-1 or ZYG-12 induces precocious FIB-1-positive nucleoli at the 4-cell stage, particularly in the EMS and P2 blastomeres. LMN-1 depletion produces a more severe phenotype combining premature assembly with impaired disassembly. Loss of the heterochromatin anchor CEC-4 alone has modest effects but strongly suppresses precocious nucleolar assembly caused by LINC complex depletion. LINC complex and lamin perturbations alter the heterogeneity of HPL-2-marked chromatin, and FIB-1 condensates occupy locally HPL-2-depleted regions. These findings identify nuclear envelope-dependent heterochromatin organization as a developmental gate for nucleolar condensate assembly.

## INTRODUCTION

The nucleolus is the largest and most prominent membraneless organelle. As the principal site of ribosome biogenesis, it coordinates the transcription, processing, and assembly of ribosomal RNA and ribosomal proteins into functional pre-ribosomal particles^1,2^. The nucleolus is organized into three immiscible liquid phases, the fibrillar center, the dense fibrillar component, and the granular component, whose layered architecture facilitates sequential steps of ribosome assembly^3^. This multiphase organization arises through liquid-liquid phase separation, driven by multivalent interactions among intrinsically disordered protein regions and RNA^1,4^. Beyond ribosome production, the nucleolus serves as a hub for stress sensing, genome organization, cell cycle regulation, and cell fate determination^2,5,6^. Nucleolar dysregulation is implicated in a diverse spectrum of human diseases, including ribosomopathies, neurodegenerative disorders, and cancer^7–11^. Mutations in nuclear lamins that cause Emery-Dreifuss muscular dystrophy (EDMD) disrupt cellular organization through a nucleolar-ribosomal axis^12^, revealing that nucleolar dysfunction can arise from defects in nuclear envelope architecture. These connections underscore the importance of understanding how this organelle is assembled, maintained, and regulated^1^.

A fundamental yet poorly understood aspect of nucleolar biology is nucleologenesis, the *de novo* assembly of nucleoli during development. In metazoan embryos, nucleoli are rebuilt after each mitotic division and, at the earliest stages, must be formed for the first time from a nucleolus-free state inherited from the mature oocyte^13,14^. In mammals, nucleolus precursor bodies appear around the time of zygotic genome activation, and their maturation into tripartite, transcriptionally active nucleoli accompanies the transition from totipotency to pluripotency^14–16^. Disrupting this process, for example through loss of RNA polymerase I activity or depletion of key nucleolar assembly factors, causes embryonic lethality^17^. These observations suggest that the timing of nucleolar assembly is not a passive consequence of increasing protein concentration but rather an actively regulated developmental transition. In *Drosophila* embryos, rRNA transcription seeds nucleolar assembly and confers its temporal and spatial precision: without rDNA, nucleolar proteins still condense, but at variable times and locations^18^, and individual nucleolar components differ in whether they assemble thermodynamically or through active, rDNA-dependent recruitment^19^. Transcription therefore provides a positive nucleating cue. It does not, however, explain why assembly is delayed. What restrains nucleolar assembly until the appropriate developmental stage remains unknown.

The *Caenorhabditis elegans* embryo offers a powerful system for addressing this question. Its invariant cell lineage, rapid cell cycles, optical transparency, and genetic tractability enable quantitative analysis of nucleolar dynamics at single-cell resolution in a living organism^20,21^. In *C. elegans*, the oocyte nucleolus dissolves prior to fertilization, and nucleoli are rebuilt during early embryogenesis^13,22^. Distinct FIB-1/DAO-5-positive nucleoli first appear at approximately the 6- to 8-cell stage, overlapping with the transition from early lineage-specific zygotic transcription to broader embryonic RNA synthesis^13,23^. Prior to the 6- to 8-cell stage, fibrillarin (FIB-1) and DAO-5/Nopp140 are diffusely distributed and sometimes form transient small nucleoplasmic foci, but rarely coalesce into recognizable nucleoli. Although nucleolar size has been linked to NCL-1 activity and nucleolar function connected to immunity and aging in *C. elegans*, why recognizable nucleoli first assemble at the 6- to 8-cell stage rather than earlier remains unknown^24–27^.

We therefore asked whether nuclear envelope architecture provides a developmental gate for nucleologenesis. The nuclear lamina, composed of intermediate filament proteins (lamins), lines the inner nuclear membrane and anchors heterochromatin at the nuclear periphery^28^. Linker of nucleoskeleton and cytoskeleton (LINC) complexes span the nuclear envelope, connecting the lamina and chromatin to the cytoskeleton in the cytoplasm^29,30^. LINC complexes consist of Sad1/UNC-84 homology (SUN) proteins in the inner nuclear membrane and Klarsicht/ANC-1/SYNE homology (KASH) proteins in the outer nuclear membrane. SUN and KASH proteins interact in the perinuclear space to form bridges across the nuclear envelope that mediate nuclear positioning, mechanical signaling, and chromatin organization^30,31^. In the early *C. elegans* embryo, LMN-1 is the sole nuclear lamin, and LINC complexes primarily consist of the SUN protein SUN-1 and the KASH protein ZYG-12^31,32^. The chromodomain protein CEC-4, an inner nuclear membrane component, tethers H3K9-methylated heterochromatin to the nuclear periphery^33^. CEC-4-mediated anchoring is genetically separable from transcriptional silencing^33^, making it possible to ask whether the spatial position of heterochromatin, independent of its transcriptional state, influences nucleolar assembly timing.

Several lines of evidence suggest that nuclear envelope architecture could influence nucleolar assembly. In mammalian cells, SUN1 regulates nucleolar morphology and rRNA synthesis, and SUN1 splice variants influence nucleolar organization in association with chromatin changes^34,35^. In *C. elegans*, the giant KASH protein ANC-1 and the nuclear lamin LMN-1 both regulate cellular biophysical properties *in vivo*, with LMN-1 additionally reported to influence cellular organization through a nucleolar-ribosomal axis^12,36^. Together with recent evidence that chromatin spatial heterogeneity can tune nuclear condensate phase behavior^37^, these observations led us to test whether LINC complexes, LMN-1, and nuclear envelope-associated heterochromatin organization gate nucleolar assembly in the early embryo. The nucleolus is also characteristically embedded within heterochromatin, and recent work has begun to define how these two condensate systems co-assemble^38,39^. In the Drosophila embryo, pericentromeric heterochromatin progressively wraps the nucleolus through a developmentally regulated hierarchy of interfacial affinities^38^. Whether the nuclear envelope and its associated chromatin-tethering machinery influence nucleolar assembly itself and whether heterochromatin organization sets the developmental timing of that assembly remains unknown.

Here, we show that the embryonic LINC complex and nuclear lamina restrain premature nucleolar assembly through heterochromatin organization at the nuclear periphery. Depletion of SUN-1 or ZYG-12 induces precocious FIB-1-positive nucleoli in 4-cell embryos, with the strongest effects in the EMS and P2 blastomeres. LMN-1 depletion produces the most severe phenotype, combining premature assembly with impaired disassembly. Loss of CEC-4 alone has modest effects, but strongly suppresses the precocious nucleolar assembly caused by LINC complex depletion. Consistently, LINC complex and lamin perturbations alter HPL-2 heterogeneity during the same cell cycles in which nucleoli assemble prematurely, and FIB-1 condensates occupy local regions with reduced HPL-2 signal. These findings identify nuclear envelope-dependent heterochromatin organization as a developmental gate for nucleolar assembly and reveal how nuclear architecture can function as a timing mechanism for condensate assembly in the early embryo.

## RESULTS

### LINC complexes restrain premature nucleolar assembly in the *C. elegans* early embryo

During nucleolar assembly, fibrillarin/FIB-1 redistributes from a diffuse nucleoplasmic pool into concentrated nucleolar condensates, providing a live readout of nucleolar assembly in early embryos^13,22,40^. We quantified this transition in living embryos expressing FIB-1::eGFP by 4D volumetric confocal imaging and segmentation, and lineage-resolved tracking of normalized maximum nuclear FIB-1::eGFP intensity (**See Methods**). In wild-type embryos, FIB-1 remained diffuse and near baseline in AB and P1 at the 2-cell stage and in most 4-cell-stage blastomeres. EMS showed only a modest, transient increase during the 4-cell stage, whereas strong FIB-1 intensity peaks appeared reproducibly in 8-cell-stage somatic blastomeres, including ABal, ABar, ABpl, ABpr, MS, E, and C. By contrast, FIB-1 in the germline precursor P3 remained largely non-condensed (**Figs. 1A and 1C**). Thus, under wild type conditions, robust FIB-1-positive nucleolar assembly is delayed until after the 4-cell stage, consistent with prior observations of early embryonic nucleologenesis in *C. elegans*^13,22,41^. Importantly, this result validates normalized maximum nuclear FIB-1::eGFP intensity as a quantitative live-cell reporter of nucleolar assembly and extends previous stage-based observations by providing lineage-resolved temporal measurements from the 2-cell through 8-cell stages.

**Figure 1.**
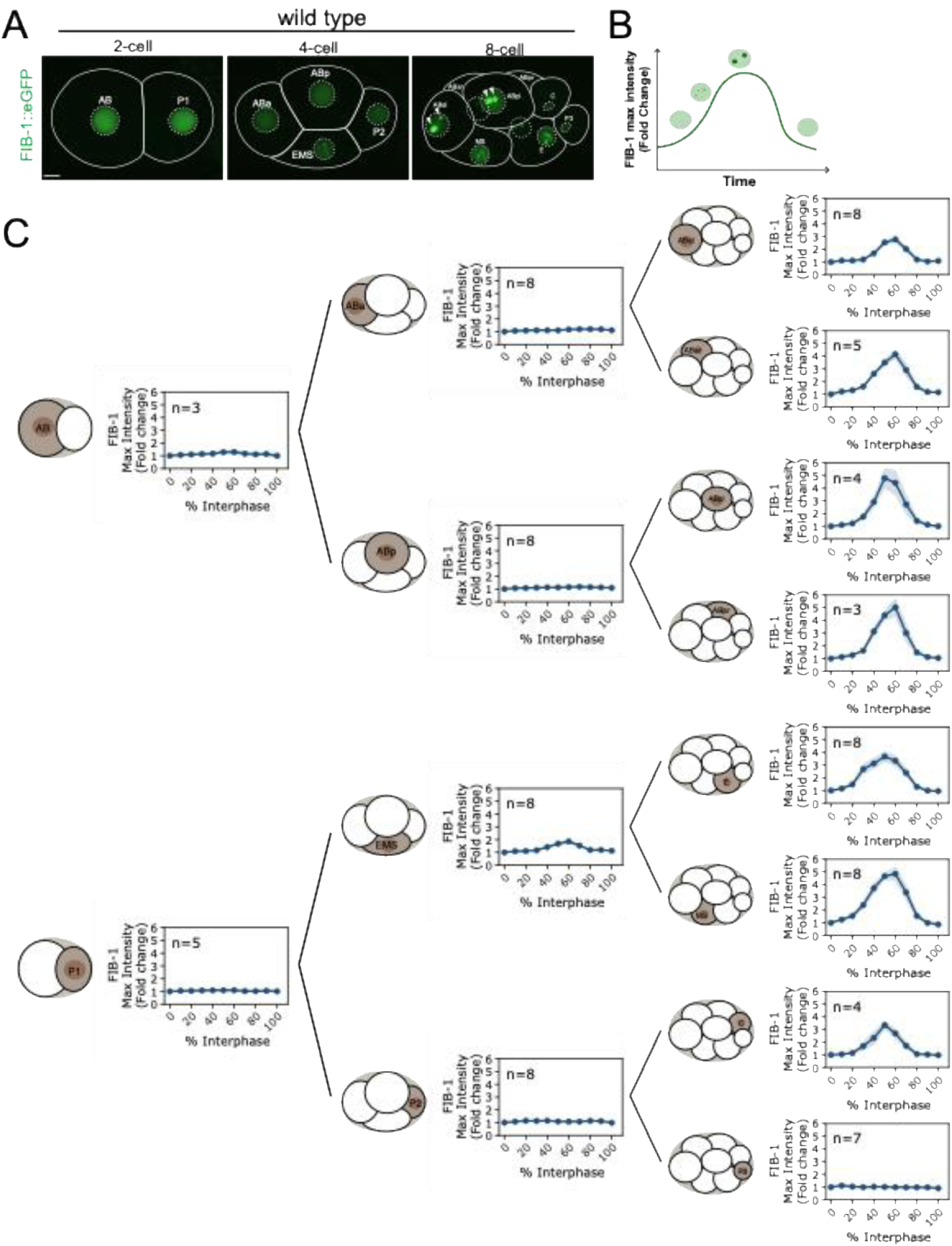
Nucleolar assembly is developmentally delayed until after the 4-cell stage. **(A)** Representative three-dimensional Imaris volume renderings, viewed along the z-axis (xy view), of wild-type 2-cell, 4-cell, and 8-cell embryos expressing FIB-1::eGFP. Cell and nuclear boundaries are indicated by solid and dashed outlines, respectively. Scale bar, 5 µm. **(B)** Schematic illustrating the use of normalized maximum nuclear FIB-1::eGFP intensity to quantify nucleolar condensation over interphase. FIB-1 intensity increases as FIB-1 transitions from a diffuse nucleoplasmic pool into concentrated nucleolar condensates and decreases upon disassembly. **(C)** Lineage-resolved trajectories of normalized maximum FIB-1::eGFP intensity in wild-type embryos. FIB-1 intensity was normalized to the first post-mitotic frame for each nucleus. Robust FIB-1 condensation is detected reproducibly in 6- to 8-cell-stage blastomeres, whereas 2- and 4-cell-stage nuclei show little or only transient accumulation. Lines indicate mean; shading indicates SEM. n values are shown in each panel.

We next asked whether the embryonic LINC complex contributes to this timing control. In early *C. elegans* embryos, the SUN protein SUN-1 and the KASH protein ZYG-12 form a nuclear-envelope-spanning LINC complex that links the nuclear interior to cytoskeletal forces (**Fig. 2A**)^31,32^. In *control(RNAi)* 4-cell embryos, FIB-1::eGFP remained largely diffuse, whereas *sun-1(RNAi)* and *zyg-12(RNAi)* embryos displayed prominent FIB-1 condensates, most visibly in EMS and P2 (**Fig. 2B**). Quantification revealed that SUN-1 or ZYG-12 depletion strongly increased FIB-1 maximum intensity during interphase of all blastomeres in the 4-cell-stage embryo, with the strongest effects in EMS and P2, whereas in the earlier AB and P1 nuclei, FIB-1 intensity remained close to baseline (**Fig. 2C**). These data indicate that the SUN-1/ZYG-12 LINC complex normally restrains FIB-1 condensation before the 6- to 8-cell stage, with EMS and P2 being most sensitive to LINC disruption.

**Figure 2.**
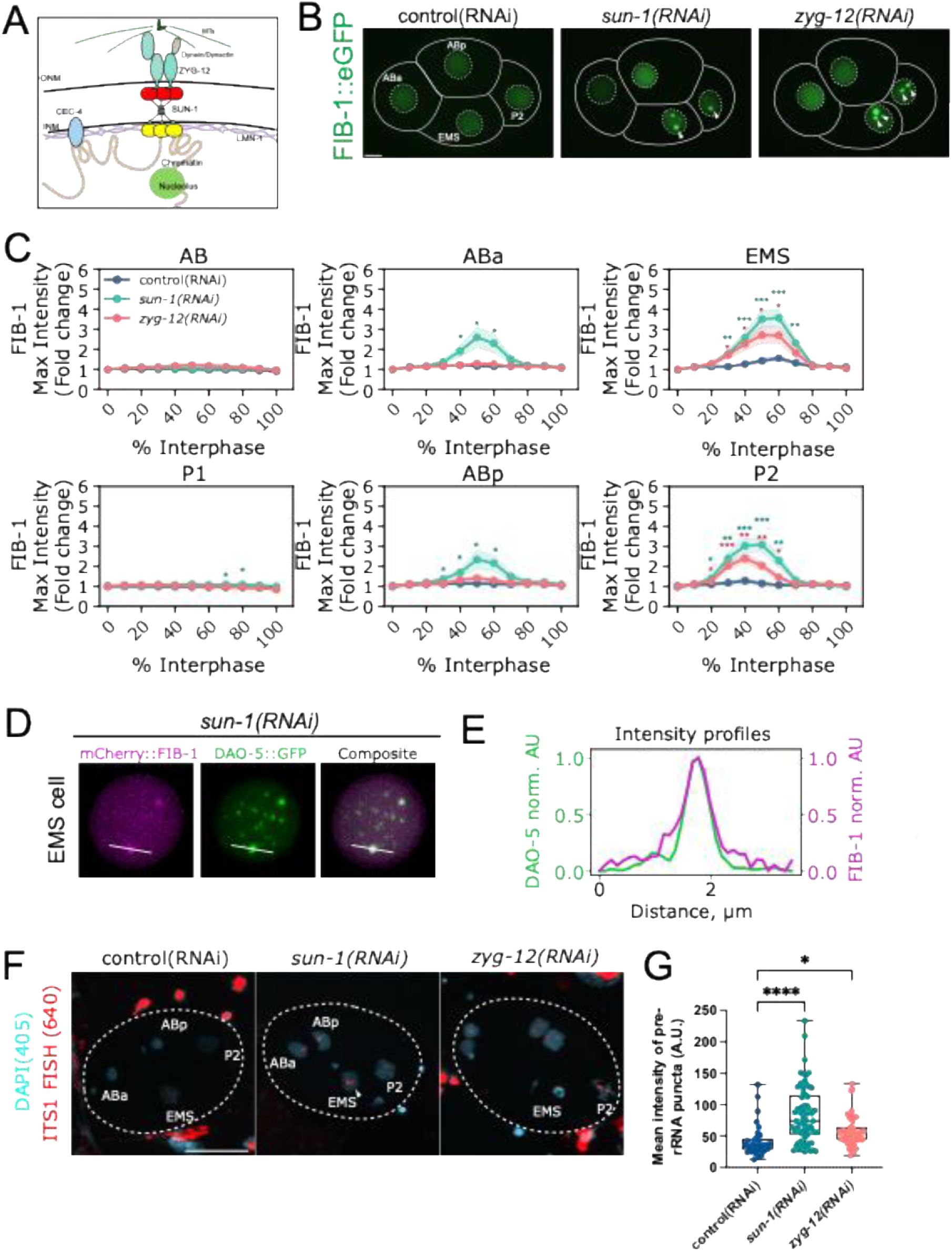
Depletion of the LINC complexes advance nucleolar assembly. **(A)** Schematic of the nuclear-envelope and chromatin-associated factors examined in this study, including the SUN-1/ZYG-12 LINC complex, LMN-1, CEC-4, chromatin, microtubule-associated forces, and the nucleolus. ONM, outer nuclear membrane; INM, inner nuclear membrane; MTs, microtubules. **(B)** Representative 4-cell embryos expressing FIB-1::eGFP in control(RNAi), *sun-1(RNAi)*, and *zyg-12(RNAi)* embryos. Arrowheads indicate precocious FIB-1 condensates. Scale bar, 10 µm. **(C)** Normalized maximum FIB-1::eGFP intensity traces for AB, P1, ABa, ABp, EMS, and P2 nuclei in control(RNAi), *sun-1(RNAi)*, and *zyg-12(RNAi)* embryos. SUN-1 or ZYG-12 depletion induces precocious FIB-1 accumulation, with the strongest effects in EMS and P2 and more modest effects in AB-lineage blastomeres. Lines indicate mean; shading indicates SEM. Asterisks mark significant differences from control(RNAi) at matched time points by two-sided Welch’s t-test; *p < 0.05, **p < 0.01, ***p < 0.001. n values for control(RNAi)/*sun-1(RNAi)/zyg-12(RNAi)*, respectively, are: AB, 7/7/7; P1, 9/5/4; ABa, 9/12/13; ABp, 9/12/13; EMS, 9/12/12; P2, 9/12/11. n indicates the number of nuclei analyzed for each blastomere and condition. **(D)** Representative EMS nucleus from a *sun-1(RNAi)* embryo co-expressing mCherry::FIB-1 and DAO-5::GFP. The white line indicates the line-scan path. **(E)** Normalized fluorescence intensity profiles measured along the line shown in (**D**), showing overlapping DAO-5::GFP and mCherry::FIB-1 intensity peaks. **(F)** Representative max-intensity projections of pre-rRNA FISH in early embryos after control(RNAi), *sun-1(RNAi)*, or *zyg-12(RNAi)* treatment. DAPI, cyan, marks DNA/nuclei imaged with the 405 nm channel; ITS1 FISH, red, marks nascent pre-rRNA signal imaged with the 640 nm channel. Dashed outlines indicate embryo boundaries, and cell identities are annotated where visible. Arrows highlight the distinct pre-rRNA signal. Scale bar, 20 µm. (**G**) Quantification of mean pre-rRNA FISH puncta intensity in the indicated RNAi conditions. Each point represents one segmented ITS1 punctum. Boxes indicate the median and interquartile range; whiskers indicate the minimum and maximum values. In total, 35 puncta from 9 *control(RNAi)* embryos, 67 puncta from 13 *sun-1(RNAi)* embryos, and 52 puncta from 8 *zyg-12(RNAi)* embryos were quantified. Statistical comparisons were performed using Brown-Forsythe and Welch ANOVA followed by Dunnett’s T3 multiple comparisons test; *p < 0.05, ****p < 0.0001.

To test whether these precociously assembled nucleolar condensates contained additional nucleolar components rather than condensates composed predominantly of FIB-1, we examined DAO-5/Nopp140, an independent nucleolar marker used previously to follow *C. elegans* nucleologenesis^13,22^. In representative *sun-1(RNAi)* EMS nucleus, DAO-5::GFP accumulated at the same ectopic focus as mCherry::FIB-1, and line-scan profile showed overlapping DAO-5 and FIB-1 intensity peaks (**Figs. 2D-E**). These observations support the interpretation that LINC complex depletion induces precocious assembly of condensates containing multiple nucleolar markers. To test whether these ectopic nucleolar assemblies were associated with pre-rRNA production, we performed hybridization chain reaction RNA fluorescence in situ hybridization (HCR RNA-FISH) targeting the ITS1 region of pre-rRNA. Compared with *control(RNAi)*, depletion of *sun-1* or *zyg-12* resulted in brighter and more prominent ITS1 FISH signals in the 4-cell stage (**Fig. 2F**). Quantification of the mean fluorescence intensity of individual ITS1-positive puncta showed increased pre-rRNA FISH signal in *sun-1(RNAi)* or *zyg-12(RNAi)* embryos (**Fig. 2G**). Together with the representative DAO-5/FIB-1 overlap, these data suggest that the precocious FIB-1 assemblies are transcriptionally active, pre-rRNA-producing nucleoli rather than FIB-1-only condensates.

### LMN-1 depletion causes precocious and persistent FIB-1 condensation

We next asked whether the nuclear lamina also contributes to nucleolar assembly timing. LMN-1 is the sole lamin in *C. elegans* and the major structural component of the nuclear lamina^42^. In *lmn-1(RNAi)* embryos, FIB-1::eGFP condensates were visible in 4-cell-stage nuclei and persisted later into the cell cycle than in control embryos (**Fig. 3A; Movie S2**). Quantification showed that LMN-1 depletion produced a gradual, sustained elevation of FIB-1 maximum intensity, with the clearest and statistically supported increases in EMS and P2 during mid-late interphase; ABa showed only a small late increase, and ABp changed little relative to control (**Fig. 3B**). Thus, LMN-1 depletion differs from SUN-1 or ZYG-12 depletion: LINC perturbation produces a pronounced mid-interphase FIB-1 condensation, whereas LMN-1 depletion causes a weaker but more persistent FIB-1 accumulation.

**Figure 3.**
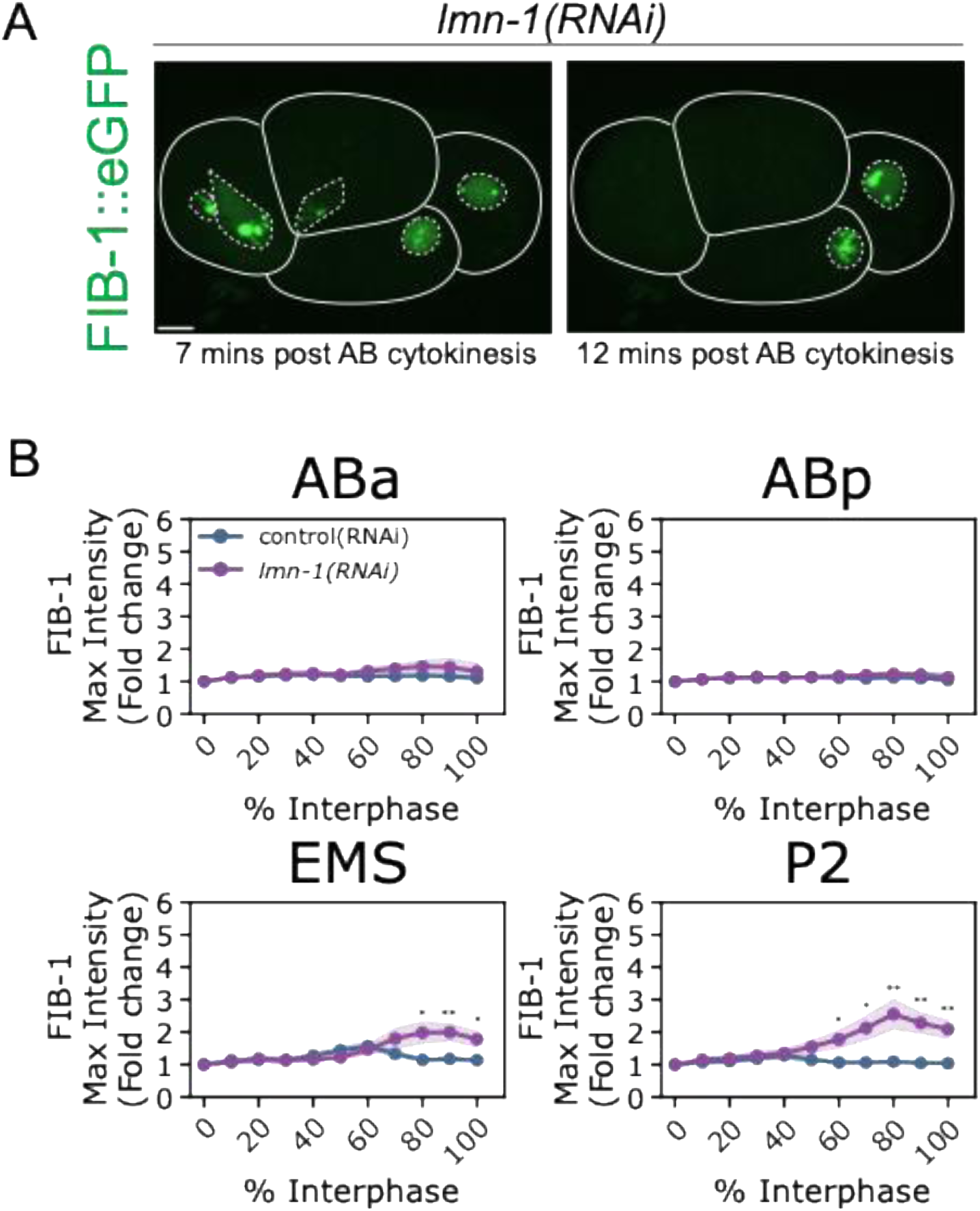
LMN-1 depletion causes both precocious nucleolar assembly and impaired disassembly. **(A)** Representative 4-cell-stage *lmn-1(RNAi)* embryos expressing FIB-1::eGFP at 7 and 12 min after AB cytokinesis. FIB-1 condensates appear prematurely and persist later into interphase. Scale bar, 10 µm. **(B)** Normalized maximum FIB-1::eGFP intensity traces for ABa, ABp, EMS, and P2 nuclei in control(RNAi) and *lmn-1(RNAi)* embryos. LMN-1 depletion increases FIB-1 accumulation most strongly in EMS and P2 and prolongs elevated FIB-1 signal compared with control embryos. Lines indicate mean; shading indicates SEM. Asterisks mark significant differences from control(RNAi) at matched time points by two-sided Welch’s t-test; *p < 0.05, **p < 0.01. n values for control(RNAi)/*lmn-1(RNAi)*, respectively, are: ABa, 9/13; ABp, 9/14; EMS, 9/14; P2, 9/14. n indicates the number of nuclei analyzed for each blastomere and condition.

Because changes in nuclear size could alter apparent FIB-1 concentration, we measured nuclear volume in EMS and P2. Nuclear volume trajectories and peak-frame nuclear volumes were not significantly altered by *sun-1(RNAi)* or *zyg-12(RNAi)*, indicating that the LINC-dependent increase in FIB-1 maximum intensity is not explained by reduced nuclear volume (**Figs. S1B-C**). *lmn-1(RNAi)* nuclei were smaller than controls during late interphase, consistent with the broader role of the lamina in nuclear structure^43^, but nuclear volume at the frame of peak FIB-1 intensity was not significantly different from control in either EMS or P2 (**Fig. S1C**). These measurements argue that the precocious FIB-1 phenotypes are not a simple consequence of nuclear volume changes, although LMN-1 depletion clearly affects nuclear architecture more broadly.

### CEC-4 is required for LINC-dependent, but not LMN-1-dependent, precocious FIB-1 Condensation

We next tested whether peripheral heterochromatin anchoring contributes to nucleolar assembly timing. CEC-4 is an inner nuclear membrane chromodomain protein that binds H3K9-methylated chromatin and anchors heterochromatin at the nuclear periphery in *C. elegans* embryos (**Fig. 2A**)^33,44^. Because CEC-4-mediated anchoring can be separated from transcriptional silencing, it provides a way to test whether heterochromatin position contributes to FIB-1 condensation^33^. If loss of peripheral anchoring were sufficient to trigger early nucleolar assembly, *cec-4(ok3124)* embryos would be expected to phenocopy LINC or lamin depletion. Instead, *cec-4(ok3124)* embryos showed no robust precocious FIB-1 accumulation in 4-cell-stage ABa, ABp, EMS, or P2 nuclei (**Figs. 4A-B**). At later stages, *cec-4(ok3124)* caused only limited changes in 8-cell-stage FIB-1 trajectories and did not broadly alter the timing of nucleolar marker accumulation (**Fig. S2A**). Thus, loss of CEC-4 alone is not sufficient to advance nucleolar assembly.

**Figure 4.**
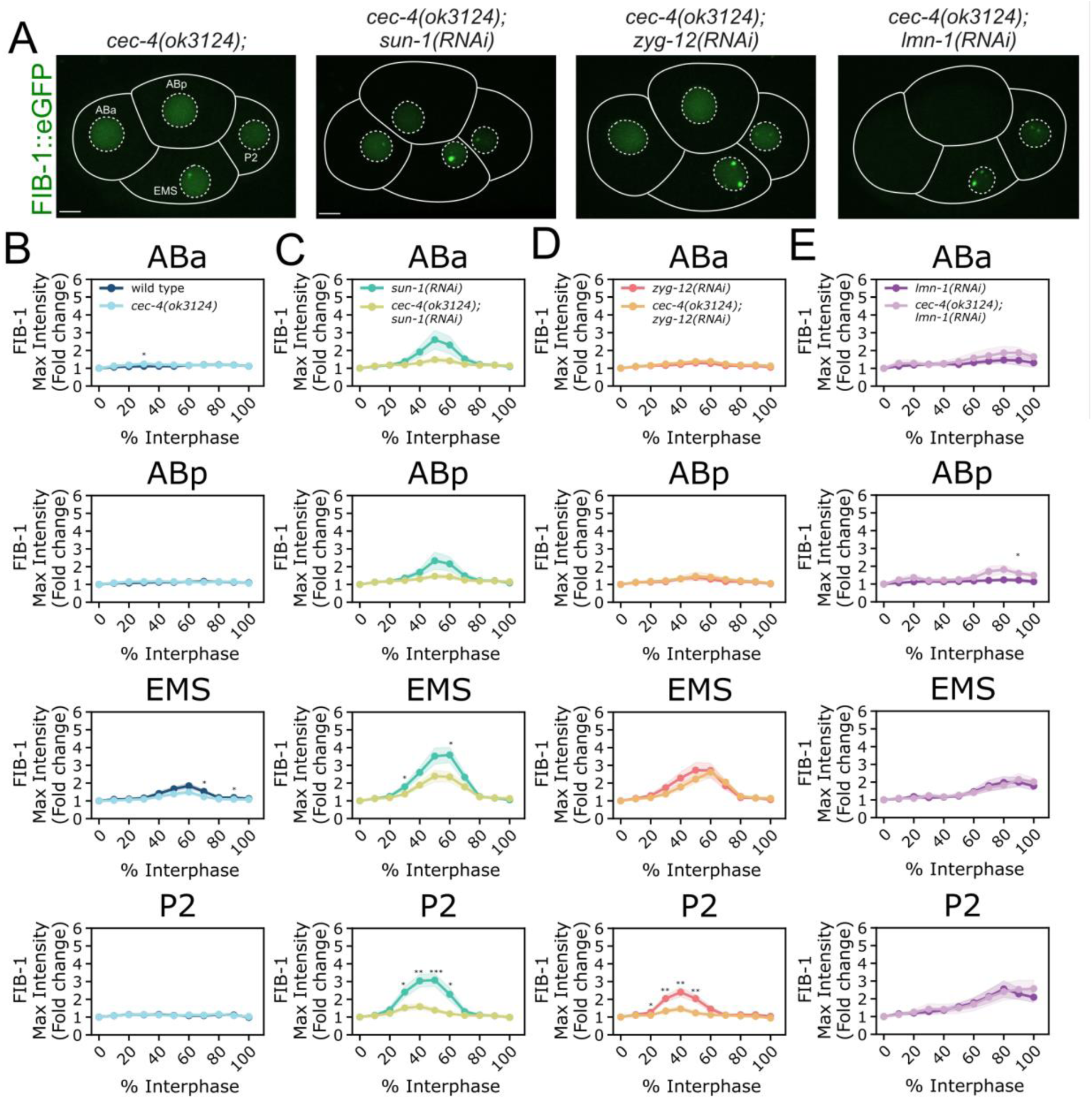
CEC-4 is required for LINC-dependent precocious nucleolar assembly. **(A)** Representative 4-cell-stage embryos expressing FIB-1::eGFP in *cec-4(ok3124)*, *cec-4(ok3124)*; *sun-1(RNAi), cec-4(ok3124)*; *zyg-12(RNAi)*, and *cec-4(ok3124)*; *lmn-1(RNAi)* backgrounds. Scale bars, 10 µm. **(B)** Normalized maximum FIB-1::eGFP intensity traces for ABa, ABp, EMS, and P2 nuclei in wild type and *cec-4(ok3124)* embryos. Loss of CEC-4 alone causes only modest changes in early FIB-1 dynamics. **(C-E)** Normalized maximum FIB-1::eGFP intensity traces comparing each single perturbation with the corresponding *cec-4* double-perturbed condition: (C) *sun-1(RNAi)* versus *cec-4(ok3124)*; *sun-1(RNAi),* (D) *zyg-12(RNAi)* versus *cec-4(ok3124); zyg-12(RNAi)*, and (E*) lmn-1(RNAi)* versus *cec-4(ok3124); lmn-1(RNAi)*. Loss of CEC-4 suppresses the precocious FIB-1 accumulation caused by SUN-1 or ZYG-12 depletion, especially in EMS and P2, but does not suppress the *lmn-1(RNAi)* phenotype. Lines indicate mean; shading indicates SEM. Asterisks mark significant differences between the paired conditions at matched time points by two-sided Welch’s t-test; *p < 0.05, **p < 0.01, ***p < 0.001. n values for the paired conditions, respectively, are: *wild type/cec-4(ok3124)*, ABa, 8/11; ABp, 8/11; EMS, 8/11; P2, 8/11; *sun-1(RNAi)/cec-4(ok3124)*; *sun-1(RNAi)*, ABa, 12/12; ABp, 12/12; EMS, 12/12; P2, 12/12; *zyg-12(RNAi)/cec-4(ok3124)*; *zyg-12(RNAi)*, ABa, 13/8; ABp, 13/7; EMS, 12/8; P2, 11/6; *lmn-1(RNAi)/cec-4(ok3124)*; *lmn-1(RNAi)*, ABa, 13/9; ABp, 14/9; EMS, 14/8; P2, 14/8. n indicates the number of nuclei analyzed for each blastomere and condition.

We then asked whether CEC-4 is required for the precocious FIB-1 condensation caused by nuclear-envelope perturbation. In *cec-4(ok3124); sun-1(RNAi)* embryos, the FIB-1 intensity peak induced by *sun-1(RNAi)* was strongly reduced in ABa, ABp, EMS, and P2 (**Fig. 4C**). In *cec-4(ok3124); zyg-12(RNAi)* embryos, FIB-1 accumulation was also attenuated relative to *zyg-12(RNAi)*, with the clearest suppression in P2 and more modest effects in other lineages (**Fig. 4D**). By contrast, *cec-4(ok3124)* did not suppress the *lmn-1(RNAi)* phenotype: FIB-1 maximum intensity remained elevated and persistent in *cec-4(ok3124); lmn-1(RNAi)* embryos (**Fig. 4E**). Measurements of individual FIB-1 condensates showed no significant matched differences in nucleolar volume or distance to the nuclear envelope after *cec-4* loss in the *sun-1(RNAi)*, *zyg-12(RNAi)*, or *lmn-1(RNAi)* backgrounds (**Fig. S2B**). Together, these genetic interactions indicate that CEC-4-dependent heterochromatin anchoring is required for the LINC-dependent precocious FIB-1 phenotype, but that LMN-1 likely influences FIB-1 condensation through broader nuclear architectural disruption.

### Nuclear-envelope perturbations alter HPL-2 heterogeneity, and FIB-1 condensates occupy locally HPL-2-depleted regions

To determine whether the nuclear-envelope perturbations that advance FIB-1 condensation also alter heterochromatin organization, we imaged mCherry::HPL-2, a *C. elegans* HP1-like protein associated with heterochromatic chromatin domains^45,46^. We quantified HPL-2 mean intensity as a bulk signal measure and the coefficient of variation (CV) as a measure of spatial heterogeneity within segmented nuclei (**Figs. 5A-B and S3A**). In control embryos, HPL-2 mean intensity increased during interphase. *sun-1(RNAi)* or *zyg-12(RNAi)* produced only modest changes compared to *control(RNAi)* in HPL-2 mean intensity, whereas *lmn-1(RNAi)* reduced the normal increase in EMS and P2 (**Fig. S3A**). Spatial heterogeneity was more strongly affected: in EMS nuclei, *sun-1(RNAi)* and *zyg-12(RNAi)* significantly increased HPL-2 CV relative to control, while *lmn-1(RNAi)* significantly decreased HPL-2 CV (**Fig. 5B**). These data show that LINC and lamin perturbations alter HPL-2 organization in distinct ways during the same interphase window in which FIB-1 condensation becomes premature or persistent.

**Figure 5.**
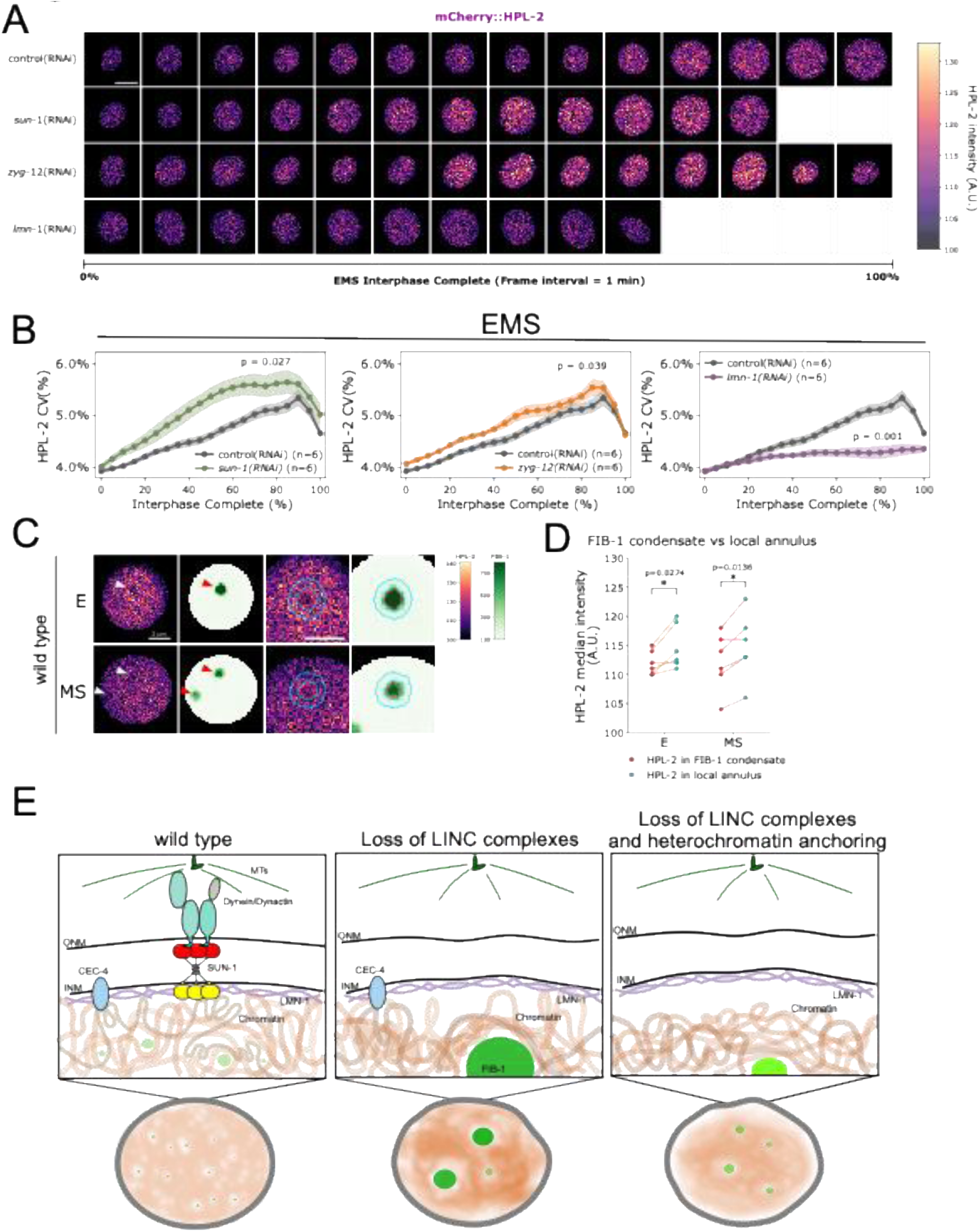
Nuclear-envelope perturbations alter HPL-2 heterogeneity, and FIB-1 condensates occupy locally HPL-2-depleted regions. **(A)** Representative time series of mCherry::HPL-2 in EMS nuclei during normalized interphase after control(RNAi), *sun-1(RNAi)*, *zyg-12(RNAi)*, or *lmn-1(RNAi)*. At each displayed time point, the single z-plane containing the maximum FIB-1 signal within the tracked EMS mask was selected independently, and the corresponding mCherry::HPL-2 signal from that same plane is shown. All panels are displayed with the same intensity scale. Frame interval, 1 min. Scale bar, 5 µm. **(B)** HPL-2 coefficient of variation in 3D across EMS interphase in the indicated RNAi conditions. CV reports spatial heterogeneity of HPL-2 signal within each nucleus. Lines indicate mean; shading indicates SEM. p values compare whole-curve AUC/100, calculated as the area under each nucleus-level interphase trajectory divided by 100 to represent the average value across normalized interphase, with control(RNAi) by two-sided Welch’s t-test. **(C)** Representative single-plane mCherry::HPL-2 and FIB-1::eGFP images of wild-type E and MS nuclei at the peak FIB-1 frame. Left to right: full HPL-2 plane, corresponding FIB-1 plane, magnified HPL-2 image, and magnified FIB-1 image. White and red arrowheads identify corresponding condensate sites in HPL-2 and FIB-1 images, respectively. Red contours indicate FIB-1 condensate cores; cyan contours indicate the corresponding local annuli. Scale bars, 2 µm. **(D)** Paired median HPL-2 intensity within each FIB-1 condensate core (red points) and its local annulus (cyan points). One condensate per embryo was selected by the strongest FIB-1 core-to-annulus enrichment; HPL-2 signal was not used for selection. Connected points indicate matched regions from the same nucleus. p values were calculated by two-sided paired t-tests (E, p = 0.0274; MS, p = 0.0136). n = 6 independent embryos per lineage. **(E)** Working model for nuclear envelope-dependent gating of nucleolar assembly in the early embryo. In wild-type 4-cell-stage embryos, SUN-1/ZYG-12, LMN-1, and CEC-4-dependent peripheral heterochromatin anchoring restrain premature FIB-1 condensation. LINC depletion increases HPL-2 heterogeneity and promotes precocious FIB-1 assembly. Loss of CEC-4 in the LINC-depleted background releases peripheral heterochromatin anchoring and suppresses premature nucleolar assembly. The local depletion of HPL-2 at FIB-1 condensates is consistent with HPL-2-marked chromatin contributing to an environment that constrains condensate formation.

We next asked whether *cec-4* loss detectably changes HPL-2 heterogeneity in the same perturbation backgrounds. In matched *cec-4(ok3124); sun-1(RNAi)* and *cec-4(ok3124); zyg-12(RNAi)* embryos, HPL-2 CV curves tended to be lower than their corresponding single-RNAi conditions in EMS or P2 lineage (**Figs. S3C-D**). By contrast, *cec-4(ok3124); lmn-1(RNAi)* embryos closely overlapped *lmn-1(RNAi)* embryos (**Fig. S3E**). Together, these matched comparisons are consistent with a model in which *cec-4* loss partially dampens the HPL-2 heterogeneity changes associated with LINC depletion, paralleling its suppression of precocious FIB-1 condensation. The close overlap between *lmn-1(RNAi) and cec-4(ok3124); lmn-1(RNAi)* embryos supports the idea that LMN-1-dependent effects on FIB-1 condensation are less dependent on CEC-4-mediated heterochromatin anchoring but likely acting through a broader or partially distinct nuclear architectural pathway.

To determine whether FIB-1 condensates are associated with local differences in HPL-2 signal, we analyzed wild-type E and MS nuclei, two lineages that robustly assemble FIB-1 condensates. At the peak FIB-1 frame for each nucleus, we selected one condensate and compared the median HPL-2 intensity within the condensate with that of its surrounding local annulus. HPL-2 signal was modestly but significantly lower within FIB-1 condensates in both E and MS (**Figs. 5C-D**). Among the remaining 8-cell-stage lineages, a significant local decrease was detected only in the ABar blastomere (**Figs. S4A-B**). However, because these measurements were obtained by 3D live-cell time-lapse imaging under acquisition conditions designed to preserve embryo viability and limit phototoxicity, the relatively weak HPL-2 signal may have reduced our sensitivity to subtle local differences. Thus, the absence of statistically significant differences in ABal, ABpl, ABpr, C, and P3 does not establish the absence of local HPL-2 depletion and may reflect either limited detection sensitivity or genuine lineage-specific variation. Together, the detectable depletion in E, MS, and ABar supports an association between FIB-1 condensates and locally reduced HPL-2 signal, consistent with a model in which HPL-2-marked chromatin helps constrain where FIB-1 condensation can occur.

## DISCUSSION

Our findings identify the nuclear envelope as an active regulator of nucleolar assembly timing in the early *C. elegans* embryo. Rather than being driven solely by increasing nucleolar protein concentration or zygotic transcription, nucleologenesis is gated by the architectural state of the nucleus. Depletion of the LINC complex components SUN-1 or ZYG-12, or of the sole lamin LMN-1, advanced FIB-1-marked nucleolar assembly to the 4-cell stage, when nucleoli are normally absent or only weakly detectable. The suppression of LINC complex-dependent precocious assembly by loss of the heterochromatin anchor CEC-4 demonstrates that this timing mechanism depends on nuclear envelope-associated heterochromatin organization. Together, these results support a model in which SUN-1/ZYG-12, LMN-1, and CEC-4-dependent peripheral heterochromatin form a regulatory circuit that gates nucleologenesis in the early embryo. More broadly, our study establishes nucleolar assembly as an *in vivo* example of developmental condensate biogenesis controlled by nuclear architecture.

### Nuclear envelope architecture links heterochromatin organization to nucleologenesis timing

Our working model **(Fig. 5E)** is that nuclear envelope-dependent heterochromatin organization determines whether the 4-cell-stage nucleus is permissive for FIB-1 condensation. In wild-type 4-cell embryos (left of Fig. 5E), SUN-1/ZYG-12 LINC complexes, LMN-1, and CEC-4-dependent peripheral heterochromatin anchoring maintain a uniform nuclear environment that restrains nucleolar assembly. In wild type E and MS nuclei, FIB-1 condensates occupy regions with modestly lower HPL-1 signal than their local surroundings, revealing a local inverse relationship. Upon depletion of LINC complexes SUN-1 or ZYG-12, the spatial heterogeneity of HPL-2 increased, creating a more permissive environment favoring condensates formation. When CEC-4 is removed in the LINC-depleted background (right of Fig 5E), heterochromatin is released from the nuclear periphery^33^, becomes more uniformly distributed, which suppresses precocious nucleolar assembly. One interpretation is that CEC-4-dependent peripheral anchoring is required for LINC complex depletion to produce the aberrant heterochromatin clustering that creates permissive zones for condensation; without anchoring, the released heterochromatin distributes more uniformly, and the permissive microenvironments are lost. Importantly, our model does not imply that chromatin depletion alone is sufficient to nucleate FIB-1 condensation. Although loss of CEC-4-mediated anchoring may reduce heterochromatin occupancy at the nuclear periphery, released chromatin is redistributed rather than leaving a stable, empty peripheral compartment. FIB-1 condensation likely requires the coincidence of a permissive low-chromatin environment^37^ with an RNA-based nucleating scaffold generated by active rDNA transcription^18,19,40,41,46,47^. In *Drosophila* embryos, rRNA transcription provides this nucleating cue and confers temporal and spatial precision on nucleolar assembly; without rDNA, nucleolar proteins still condense, but sporadically, at variable times and locations^18^. Our ITS1 FISH data show that precocious nucleoli in LINC complex-depleted embryos are transcriptionally active, indicating that the nucleating cue is available before the 6- to 8-cell stage and that nuclear envelope-dependent heterochromatin organization, rather than transcription, sets the timing of assembly. Thus, peripheral regions vacated by heterochromatin may lack the molecular cues required for nucleolar assembly. Consistent with this distinction, HPL-2 signal is locally reduced within FIB-1 condensates, but not every HPL-2-depleted region forms a condensate. We therefore interpret low chromatin occupancy as permissive, but not sufficient, for FIB-1 condensation.

This model is consistent with prior work showing that nuclear envelope proteins can regulate nucleolar morphology and activity. In mammalian cells, SUN1 depletion alters nucleolar size and rRNA synthesis, and SUN1 splice variants regulate nucleolar morphology in association with changes in chromatin organization^34,35^. Lamins also influence nucleolar physiology, although the direction of the effect depends on context. Lamin B1 depletion disrupts nucleolar organization and reduces RNA synthesis in HeLa cells, whereas progerin expression or lamin A/C depletion enlarges nucleoli and increases nucleolar rRNA synthesis in human fibroblasts^48,49^. Our study adds a developmental dimension to this literature by showing that SUN-1/ZYG-12 and LMN-1 regulate both nucleolar morphology and the timing of nucleolar assembly *in vivo*.

### Lineage-specific penetrance of nucleolar timing phenotypes

The precocious nucleolar assembly phenotypes described here were most frequent in EMS and P2, with more modest effects in ABa and ABp. EMS is the first blastomere to initiate robust zygotic transcription, and P2 inherits distinct germline determinants that influence chromatin state^23,50,51^. We posit that these lineage-specific transcriptional and chromatin landscapes create different proximities to the nucleolar assembly threshold, such that a perturbation to heterochromatin organization pushes EMS and P2 over the threshold before AB-derived cells.

### LINC complexes and LMN-1 act at distinct architectural levels

The distinct phenotypes of LINC complex or lamin depletion suggest that these nuclear envelope components regulate nucleolar timing through partially overlapping but non-identical mechanisms. SUN-1 and ZYG-12 likely regulate specific chromatin-associated or force-coupled inputs into the nucleolar assembly gate. In *C. elegans*, SUN-1 colocalizes with LMN-1 and binds lamin *in vitro*, although its interphase nuclear-envelope localization is not entirely lamin-dependent^52,53^. SUN-1 and ZYG-12 also organize chromosome behavior at the nuclear envelope. SUN-1/ZYG-12-associated chromosome attachment sites connect meiotic chromosome pairing centers to microtubule- and dynein-dependent forces to promote homolog pairing and synapsis^54–57^, establishing the precedent that SUN-1/ZYG-12 can influence chromosome organization through nuclear-envelope-associated tethering and force transmission. By contrast, LMN-1 depletion compromises a broader nuclear scaffold. This explains why LMN-1 loss produces a more severe phenotype, combining premature FIB-1 assembly with impaired disassembly. In LINC knockdown embryos, precocious FIB-1 condensates dissolve prior to mitosis, consistent with the global nuclear reorganization that accompanies nuclear envelope breakdown. However, in *lmn-1(RNAi)* embryos, elevated FIB-1 signal persists until loss of nuclear integrity, suggesting that lamin functions independently of the LINC complex-dependent heterochromatin gating mechanism.

Lamin depletion also impairs nucleolar disassembly, a phenotype not seen in embryos where the LINC complex was disrupted. In our model, the persistent condensates may reflect an altered material state of the nucleolus in lamin-depleted nuclei, where nucleolar condensates become more solid-like and resistant to the forces that normally drive mitotic dissolution. The lack of suppression of the *lmn-1* phenotype by CEC-4 loss further supports the view that LMN-1 depletion disrupts a broader nuclear scaffold that integrates LINC-dependent inputs, peripheral heterochromatin organization, and potentially the rDNA–nucleolar environment. This distinction is consistent with recent *C. elegans* studies showing that ribosome concentration and the giant KASH protein ANC-1 shape cytoplasmic material properties *in vivo*, and that EDMD-associated *lmn-1* variants disrupt macromolecular crowding through a nucleolar-ribosomal axis^12,36^. Those studies linked nuclear envelope proteins, ribosomes, and nucleolar physiology to cell-scale biophysical organization, but left open how nuclear envelope architecture communicates with the nucleolus. The present study addresses this gap by showing that SUN-1/ZYG-12 and LMN-1 regulate nucleolar assembly timing through heterochromatin organization.

### Heterochromatin organization and condensate permissiveness

Chromatin spatial heterogeneity has been shown to create permissive or non-permissive microenvironments that modulate condensate growth, mobility, and phase behavior in cultured mammalian cells^37^. Our findings suggest that this biophysical principle may contribute to the developmental timing of nucleolar assembly. In several wild type lineages, HPL-2 signal was modestly reduced within FIB-1 condensates relative to their local surroundings, revealing an inverse spatial association between HPL-2-marked chromatin and FIB-1 condensation (**Figs. 5C-D and S4**). We therefore propose that the permissiveness of the nuclear environment for condensate formation is actively regulated through LINC complexes, nuclear lamins, and CEC-4-dependent heterochromatin anchoring. Although LINC complex depletion increased HPL-2 heterogeneity and lamin depletion decreased it, both promoted precocious nucleolar assembly. These contrasting effects may reflect distinct mechanisms: LINC complex depletion may over-cluster heterochromatin, creating local chromatin-depleted zones permissive for condensation, whereas LMN-1 depletion may more broadly disrupt nuclear organization and weaken chromatin-mediated constraints on nucleolar assembly. Both perturbations therefore shift the nucleus away from the non-permissive wild-type state. This interpretation is consistent with findings in cultured mammalian cells showing that both excessive clustering and excessive homogenization of chromatin can alter condensate phase behavior^37^. However, the sensitivity limits of live imaging currently preclude determining whether local HPL-2 depletion is a general feature of FIB-1 condensates or varies among lineages. Future work using more sensitive imaging approaches will be needed to resolve this question and determine how chromatin architecture intersects with rDNA positioning, nascent rRNA transcription, and nuclear biophysical properties to control nucleologenesis.

### Relationship to nucleolar-heterochromatin co-assembly

Heterochromatin is a prominent feature of the nucleolar periphery across diverse eukaryotes, and recent work has begun to define how these two condensate systems co-organize^6,38,39^. Rajshekar et al. showed that pericentromeric heterochromatin progressively surrounds the nucleolus over Drosophila embryogenesis, governed by a hierarchy of interfacial tensions among heterochromatin, nucleolar subcompartments, and a dual-affinity linker, the DEAD-box helicase Pitchoune (DDX18)^38^. Our observations are concordant with theirs in showing that, in specific lineages, heterochromatin and nucleolar phases are locally anti-correlated: FIB-1 condensates in E and MS occupy different microenvironments in which HPL-2 is locally depleted relative to the surrounding nucleoplasm (**Figs. 5C-D**), although this local exclusion is not uniformly significant across all blastomeres (**Fig. S4**). The two studies nonetheless address distinct regulatory layers. Rajshekar et al. define how an existing nucleolus organizes the heterochromatin around it, with the causal arrow running from the nucleolus to heterochromatin^38^. Falahati et al. (2016) define a second arrow, running from transcription to the nucleolus: rRNA seeds assembly and sets its precision^18^. Our data instead identify an upstream determinant of nucleolar assembly itself: LINC complex- and lamin-dependent organization of heterochromatin sets the developmental timing at which nucleoli are permitted to assemble. Depletion of SUN-1, ZYG-12, or LMN-1 reorganizes heterochromatin and triggers precocious nucleolar assembly, a phenotype with no counterpart in a wetting-hierarchy framework that presumes a formed nucleolus. Thus, whereas affinity hierarchies explain the steady-state geometry of co-assembled condensates, and rRNA transcription explains when assembly can be nucleated, our results reveal that the nuclear envelope acts as a temporal gate on condensate biogenesis during development, coupling architectural inputs at the nuclear periphery to the chromatin landscape that licenses phase separation.

### Implications and future directions

It remains important to determine what nucleolar assembly contributes to post embryonic fitness. Because SUN-1, ZYG-12, and LMN-1 perturbations also affect mitosis, nuclear integrity, and cell-cycle progression^31,32,42^, downstream developmental consequences cannot yet be attributed to nucleolar timing specifically. Identifying perturbations that shift nucleologenesis without broadly disrupting embryogenesis will be necessary to test whether premature or delayed nucleolar assembly affects embryonic patterning, viability, or postembryonic development. More broadly, our study demonstrates that developmental control of condensate biogenesis is shaped not only by local molecular concentration or transcriptional activation, but by nuclear envelope-dependent chromatin organization. This framework may help explain why nuclear envelope defects and nucleolar dysfunction are coupled in laminopathies, aging, and cancer^12,26,49,58^.

## METHODS

### *C. elegans* strains and maintenance. UD87

(unc-119(ed3) III; knuSi221 [fib-1p::fib-1(genomic)::eGFP::fib-1 3’UTR + unc-119(+)] II; ndc-1::mRuby V); **UD930** (*knuSi221 [fib-1p::fib-1(genomic)::eGFP::fib-1 3’UTR + unc-119(+)] II; cec-4(ok3124) IV; ndc-1::mRuby V*); **UD1021** (*unc-119(ed3) III; knuSi221 [fib-1p::fib-1(genomic)::eGFP::fib-1 3’UTR + unc-119(+)] II; wjIs104 [pie-1p::mCherry::hpl-2 + unc-119(+)]*); **UD1022** (*ptnsIs050 [DAO-5::GFP] I; unc-119(ed3) III; wjIs103 [pie-1p::mCherry::fib-1 + unc-119(+)]*); **UD1142** (*unc-119(ed3) III; knuSi221 [fib-1p::fib-1(genomic)::eGFP::fib-1 3’UTR + unc-119(+)] II; wjIs104 [pie-1p::mCherry::hpl-2 + unc-119(+)]; cec-4(ok3124) IV*). The *knuSi221 [fib-1p::fib-1(genomic)::eGFP::fib-1 3′UTR + unc-119(+)]* reporter was originally described by Allen et al^59^. and was obtained from CGC for this study, and *cec-4(ok3124)* was previously characterized by Gonzalez-Sandoval et al.^33^. *wjIs103 [pie-1p::mCherry::fib-1]* and *wjIs104 [pie-1p::mCherry::hpl-2]* were originally generated in the Arai et al^46^. Worms were maintained on standard nematode growth medium (NGM) agar plates seeded with *Escherichia coli* OP50 at 20° C under standard conditions.

### RNAi by feeding

Gene knockdown was performed using RNAi by feeding using standard methodology. Bacterial strains harboring gene-specific RNAi clones were obtained from the Ahringer RNAi library^60^ (Source Bioscience, Nottingham, UK) and verified by Sanger sequencing prior to use. Individual colonies were inoculated into 2 mL LB medium supplemented with 100 µg/mL carbenicillin in 15 mL snap-cap tubes and grown overnight at 37° C with shaking. Overnight cultures were diluted 1:40 into 2 mL fresh LB containing 100 µg/mL carbenicillin and grown for 3 h at 37° C. For dsRNA induction, 2 mL pre-warmed LB containing 100 µg/mL carbenicillin and 1 mM isopropyl β-D-1-thiogalactopyranoside (IPTG) was added to each culture to achieve a final IPTG concentration of 0.5 mM, and cultures were incubated for an additional 3∼4 h at 37° C. Induced bacterial cultures were seeded onto NGM agar plates containing 25 µg/mL carbenicillin and 1 mM IPTG and allowed to dry at room temperature. Synchronized L2– L3 larvae were transferred onto RNAi plates and maintained at 20° C for 48 h until they reached the young adult stage.

### *C. elegans* embryo dissection and sample preparation

L4-stage hermaphrodites were confirmed 1 day prior to imaging and maintained at 20 °C to obtain young adults with early embryos. For embryo dissections, 6-8 young adults were transferred into an 8 µL drop of egg buffer (4 mM HEPES (pH 7.4), 94 mM NaCl, 3.2 mM KCl, 2.7 mM CaCl₂, and 2.7 mM MgCl₂) on a 22 × 22 mm coverslip. Embryos were released by cutting the worms with the edge of a 25-gauge needle. The coverslip was then inverted onto a 2% agar pad on a glass slide. The edges were immediately sealed with melted VALAP (1:1:1 Vaseline: lanolin: paraffin) to prevent evaporation during time-lapse imaging. The slides were then mounted on the microscope immediately.

### 4D volumetric time-lapse confocal microscopy

Embryos were imaged on an Andor BC43 microscope in confocal mode using a 60x/1.42 NA oil-immersion objective. For GFP tagged proteins, including FIB-1::eGFP and DAO-5::GFP, samples were excited with a 488 nm laser at 5% laser power with a 500 ms exposure per z-plane. For mCherry::HPL-2, samples were excited with a 561 nm laser at 5% laser power with a 500 ms exposure per z-plane. For dual-color DAO-5::GFP and mCherry::FIB-1 imaging, DAO-5::GFP was acquired with 488 nm excitation at 5% laser power, and mCherry::FIB-1 was acquired with 561 nm excitation at 30% laser power, both with 500 ms exposure per z-plane. Z-stacks spanning the full embryo volume were acquired with a 1 um step size. Time-lapse series were collected at 60 s intervals for 4D imaging in x, y, z, and time.

### Quantitative imaging analysis

4D time-lapse datasets were analyzed using Imaris 10.2. Nuclei were segmented using surfaces module with machine-learning pixel classification on the FIB-1::eGFP channel. Foreground and background classes were trained using manual brush annotations, and the resulting classifier was applied to the 488 nm channel to generate nuclear surface masks. Surface smoothing was enabled with a grain size of 0.6 µm, background subtraction and region growing were disabled, and segmented nuclei were tracked in Imaris using the autoregressive motion algorithm and manually assigned to embryonic lineages. For each nucleus, voxel intensity of FIB-1::eGFP or mCherry::HPL-2 within the segmented nuclear volume was quantified at every time point. For each cell cycle, 0% interphase or t=0 was defined as the first frame in which a newly reformed nucleus became detectable after completion of mitosis, and the completion of interphase or the end of the trace was defined as the last frame the nuclei is detected before subsequent mitosis, when nuclear envelope breakdown rendered FIB-1::eGFP marked nuclei no longer distinguishable. For each nucleus, the maximum FIB-1::eGFP intensity at each time point was divided by its value at t=0, yielding a fold-change trajectory over interphase. For line-scan analysis, fluorescence intensity profiles were measured along the line from both channels 488 nm and 561 nm maximum z-projected images. For normalized plots, each channel profile was independently normalized using percentile min-max scaling, with the 5th percentile set to 0 and the 99th percentile set to 1, followed by clipping to the 0-1 range.

### HPL-2 heterogeneity analysis

Spatial heterogeneity of mCherry::HPL-2 was quantified within each segmented three-dimensional nuclear volume as the coefficient of variation (standard deviation divided by mean voxel intensity, expressed as a percentage) at each time point. Individual nuclear trajectories were rescaled from 0% to 100% of tracked interphase before averaging across nuclei. For the representative time series in Fig. 5A and Fig. S3B, mCherry::HPL-2 was displayed from the single z-plane containing the maximum FIB-1::eGFP signal within the tracked nuclear mask at each time point; the corresponding HPL-2 signal from the same plane is shown.

### Local HPL-2 analysis at FIB-1 condensates

Wild type E and MS nuclei were analyzed at the frame of peak FIB-1::eGFP signal. For each z-plane, FIB-1 condensate cores were identified within the two-dimensional nuclear mask as connected pixels with intensities at or above the 98th percentile of nuclear FIB-1 values. This percentile-based threshold restricts segmentation to the brightest nuclear FIB-1 signal and reduces the inclusion of background intensity fluctuations. For each core, a local annulus extending more than one and up to five pixels from the core boundary was restricted to the nuclear mask; other condensate cores and their adjacent guard pixels were excluded. One condensate per lineage was quantified based on the FIB-1 core-to-annulus enrichment, without reference to HPL-2 intensity. Median mCherry::HPL-2 intensity was measured within the condensate core and its corresponding annulus. Paired core and annulus intensities were compared separately for E and MS using two-sided paired t-tests.

### HCR Gold RNA-FISH for pre-rRNA ITS1

Pre-rRNA ITS1 was detected in *C. elegans* embryos using HCR Gold RNA-FISH (Molecular Instruments) modified from the manufacturer’s sample-on-slide protocol. Clean glass slides were coated the day before use with 100-200 µL 0.01% poly-L-lysine over the sample area for 10 min at room temperature, drained, air-dried completely in a dust-free slide box, and stored overnight. On the day of fixation, 20-30 gravid adults were transferred in 5-6 µL M9 onto the coated area, then covered with a coverslip, and gently compressed the coverslip with tweezers to release embryos under the dissecting microscope. Slides were plunged into liquid nitrogen for 10-15 s, the coverslip was removed by freeze-crack, and the slide was immediately immersed in prechilled 100% methanol (−20 °C) for 5 s. Methanol was drained by blotting the slide edge on Kimwipes, and embryos were fixed with 1 mL 4% paraformaldehyde in PBS for 10 min at room temperature in a humidified chamber. After fixation, samples were equilibrated for 5 min at room temperature in 200 µL of a 1:1 mixture of HCR HiFi Probe Hybridization Buffer and PBST (PBS with 0.1% Tween-20), then prehybridized in 200 µL HCR HiFi Probe Hybridization Buffer for 30 min at 37 °C. A custom HCR HiFi probe set (standard scale; Molecular Instruments; for amplifier X3) targeted a 464 nt ITS1 sequence from the *C. elegans* N2 rRNA/45S precursor (NCBI accession MN519140.1, bases 2688-3151), enabling detection of pre-rRNA processing intermediates. For each slide, probe solution was prepared as 2 µL HCR HiFi Probe in 98 µL HCR HiFi Probe Hybridization Buffer; 50-100 µL was applied to the embryo area, covered with a clean coverslip, and hybridized overnight at 37 °C in a humidified chamber.

On the next day, coverslips were floated off in prewarmed 1x HCR HiFi Probe Wash Buffer, and slides were washed four times for 15 min at 37 °C in 1x HCR HiFi Probe Wash Buffer followed by three 5 min washes in 5x SSCT (5x SSC with 0.1% Tween-20). Samples were preamplified for 30 min at room temperature in HCR Gold Amplifier Buffer. HCR Gold amplifier X3-647 hairpins h1 and h2 were snap-cooled separately by heating at 95 °C for 90 s and cooling to room temperature in the dark for 30 min, then 2 µL h1, 2 µL h2, and 96 µL HCR Gold Amplifier Buffer were combined and 100 µl were applied per slide. Amplifier solution was applied to each sample under a coverslip for 4 h at room temperature in a humidified chamber in the dark. Coverslips were floated off in room-temperature HCR Gold Amplifier Wash Buffer, and slides were washed in HCR Gold Amplifier Wash Buffer twice for 5 min, twice for 30 min, and once for 5 min. Embryos were counterstained with DAPI for 15 min at room temperature in the dark, briefly washed twice in 5x SSCT, mounted wet in ProLong Diamond Antifade Mountant (Invitrogen), and sealed with VALAP. Mounted HCR RNA-FISH samples were imaged on an Andor BC43 microscope in confocal mode. DAPI-stained nuclei were acquired using 405 nm excitation at 5% laser power with a 100 ms exposure per z-plane. The HCR-amplified pre-rRNA ITS1 signal generated by the X3-647 hairpins was acquired using 640 nm excitation at 5% laser power with a 100 ms exposure per z-plane. Three-dimensional z-stacks spanning 24 µm along the z-axis were acquired with a 0.2 µm step size. Pre-rRNA ITS1 signal was then quantified from confocal z-stacks of HCR RNA-FISH samples. For each embryo, maximum-intensity projections were generated from the 640 nm ITS1 FISH channel, and ITS1-positive puncta were manually segmented in Fiji/ImageJ. The mean fluorescence intensity within each segmented punctum was measured from the ITS1 channel. In total, 35 puncta from 9 control(RNAi) embryos, 67 puncta from 13 *sun-1(RNAi)* embryos, and 52 puncta from 8 *zyg-12(RNAi)* embryos were quantified. Puncta measurements were pooled by condition for visualization and statistical analysis.

### Statistical analysis

Statistical analyses were performed as described in the figure legends. Unless otherwise indicated, comparisons between two groups were performed using two-sided Welch’s t-tests, and paired comparisons were performed using two-sided paired t-tests. For multiple group comparisons with unequal variances, Brown-Forsythe and Welch ANOVA followed by Dunnett’s T3 multiple comparisons test was used. For time-course data, comparisons were performed at matched time points or on whole-curve summary values as indicated in the figure legends. The definition of n, error bars, boxplot elements, and p-value thresholds are specified in each figure legend.

## DATA AVAILABILITY

All *C. elegans* strains listed in the method section are available from the corresponding authors (Dan Starr; G.W.G. Luxton) upon request. Requests from academic researchers will be fulfilled promptly, with shipping costs covered by the requesting laboratory. Requests from for-profit organizations will be handled through the UC Davis Technology Transfer Office using a standard Material Transfer Agreement. In addition, frequently requested strains will be deposited in the Caenorhabditis Genetics Center (CGC), funded by the NIH Office of Research Infrastructure Programs (P40 OD010440), where they will be publicly available.

## ACKNOWLEDGEMENTS

We thank members of the Starr-Luxton and Weber labs for helpful discussions; Dr. Thomas Wilkop and the UC Davis MCB Light Imaging Facility for imaging support; WormBase for community resources; and the Caenorhabditis Genetics Center (CGC), which is funded by the NIH Office of Research Infrastructure Programs (P40 OD010440). We also thank Dr. Akatsuki Kimura for kindly providing the original strains carrying the wjIs104 [pie-1p::mCherry::hpl-2 + unc-119(+)] and wjIs103 [pie-1p::mCherry::fib-1 + unc-119(+)] alleles.

## AUTHOR CONTRIBUTIONS

Conceptualization: XD, SCW, DAS, GWGL

Methodology: XD, GW Software: XD, GW, DE

Validation: XD, GW Formal analysis: XD, GW

Resources: XD, DAS, GWGL Data curation: XD

Investigation: XD, GW, DE, ML, SZ, MC

Visualization: XD, GW

Supervision: DAS, GWGL

Project administration: XD, GWGL

Funding acquisition: DAS, GWGL

Writing-original draft: XD, DAS, GWGL

Writing-review & editing: XD, SCW, DAS, GWGL

## SUPPLEMENTAL FIGURES

**Figure S1.**
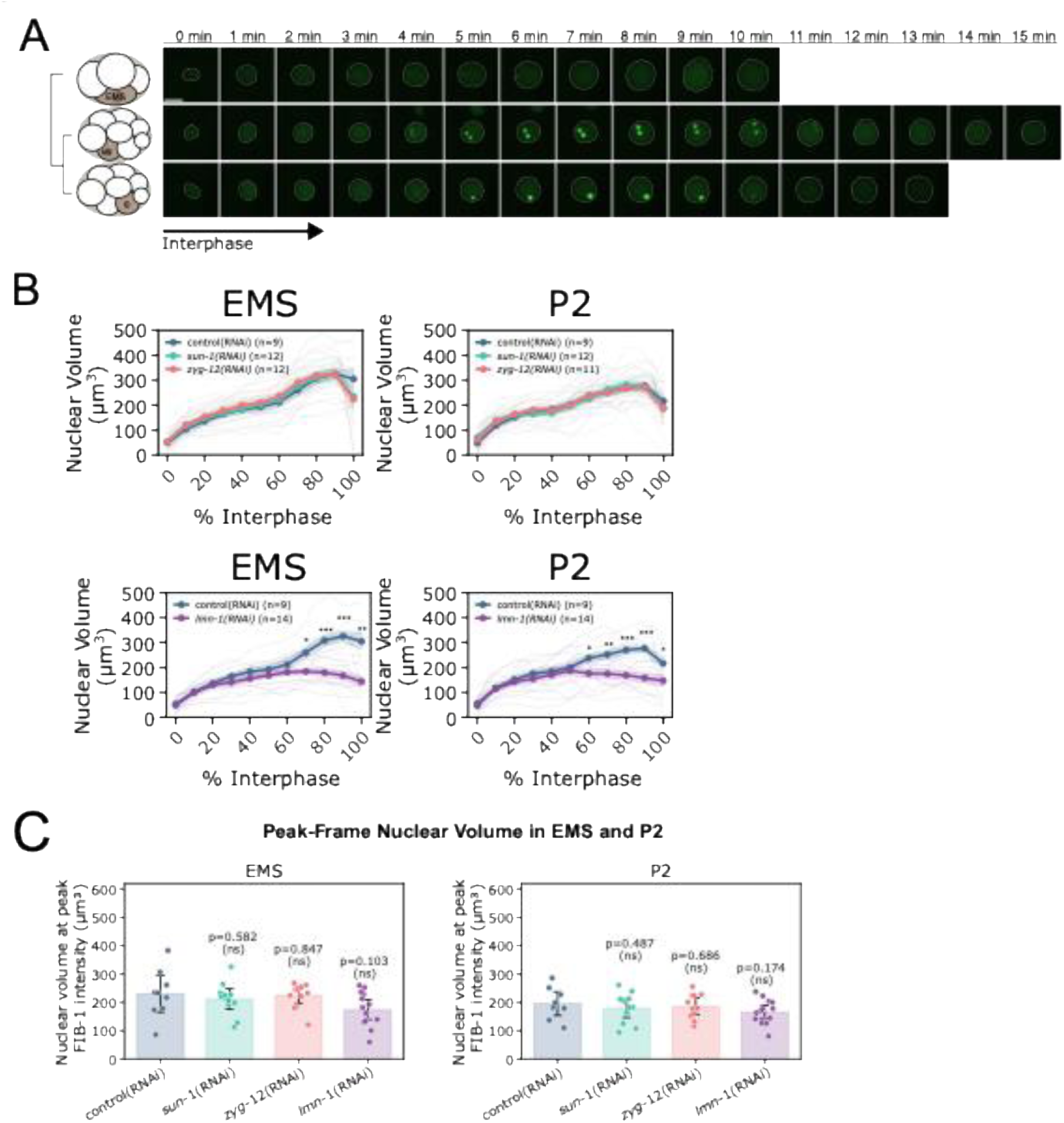
Nuclear volume changes do not account for precocious nucleolar assembly in *sun-1(RNAi)* or *zyg-12(RNAi)* embryos. **(A)** Representative time-course images of FIB-1 signal in the EMS, MS, and E nuclei during interphase. Images are maximum-intensity projections; dashed outlines indicate the segmented nuclear boundary. Time is shown in minutes relative to the start of the displayed interphase interval. All images are displayed with the same green LUT and identical intensity range for comparison. Scale bar, 5 µm. **(B)** Nuclear volume trajectories across normalized interphase in EMS and P2 nuclei after control(RNAi), *sun-1(RNAi)*, *zyg-12(RNAi)*, or *lmn-1(RNAi)*. Lines indicate mean; shading indicates SEM. Asterisks mark significant differences from control(RNAi) at matched time points by two-sided Welch’s t-test; *p < 0.05, **p < 0.01, ***p < 0.001. n values are indicated in each plot. **(C)** Nuclear volume at the frame of peak FIB-1 intensity in EMS and P2 nuclei. Each point represents one nucleus; bars indicate mean ± 95% CI. Peak frame was defined as the frame with maximal FIB-1 intensity for each nucleus. p values compare each RNAi condition with control(RNAi) by Welch’s t-test; ns, not significant. n values for control(RNAi)/*sun-1(RNAi)/zyg-12(RNAi)/lmn-1(RNAi)*, respectively, are: EMS, 9/12/12/14; P2, 9/12/11/14. n indicates the number of nuclei analyzed for each blastomere and condition.

**Figure S2.**
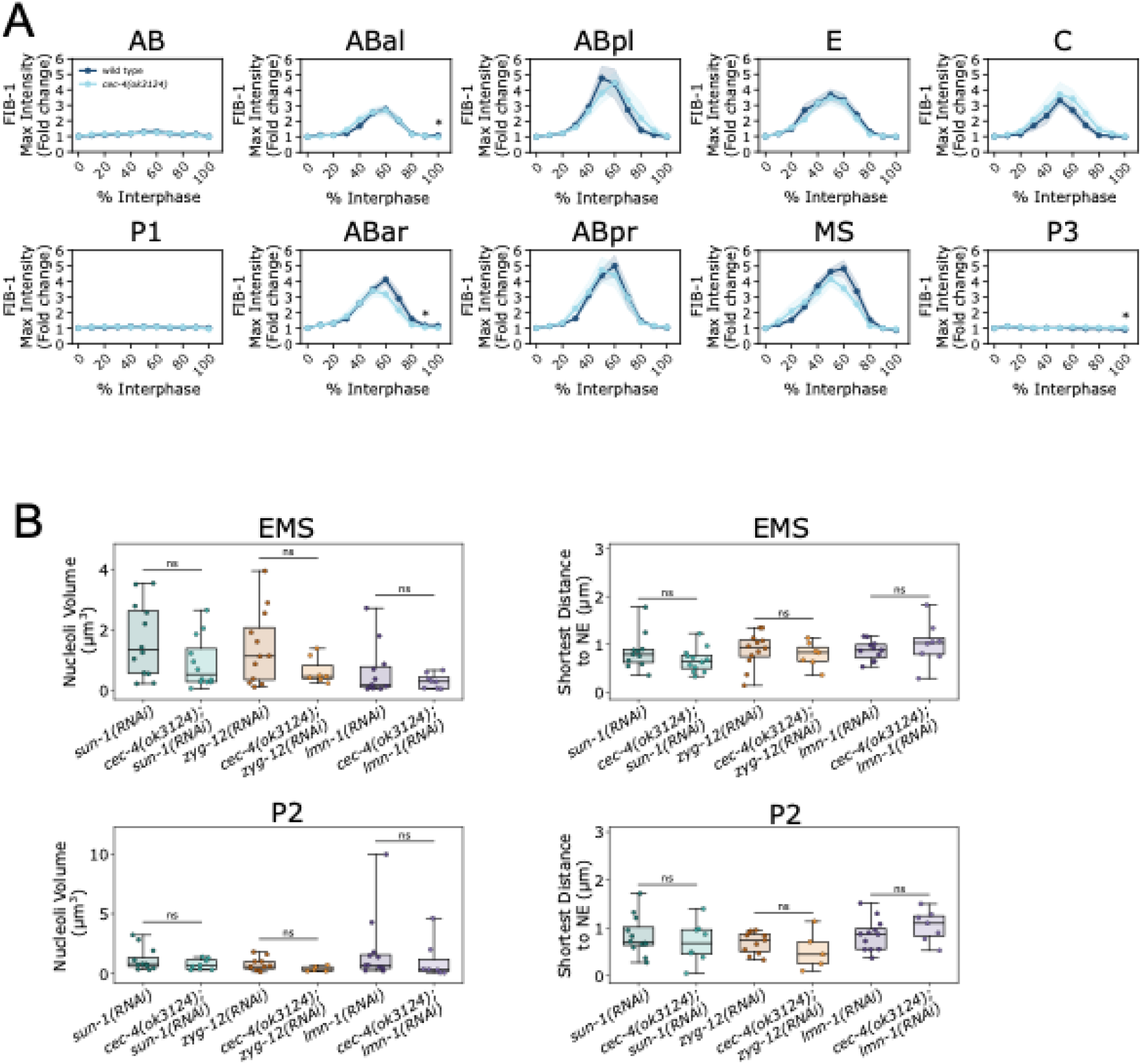
CEC-4 loss has limited effects on 8-cell-stage nucleologenesis and does not alter nucleolar size or position in double-perturbed embryos. **(A)** Normalized maximum FIB-1::eGFP intensity traces for AB, P1, and 8-cell-stage lineages in wild-type and *cec-4(ok3124)* embryos. FIB-1 intensity was normalized to the first tracked frame and plotted across normalized interphase. Lines indicate mean; shading indicates SEM. Asterisks mark significant differences between wild type and *cec-4(ok3124)* at matched time points by two-sided Welch’s t-test; *p < 0.05. n values for wild type/*cec-4(ok3124)*, respectively, are: AB, 3/5; P1, 5/6; ABal, 8/11; ABar, 5/6; ABpl, 4/5; ABpr, 3/6; MS, 8/10; E, 8/11; C, 4/6; P3, 7/5. n indicates the number of nuclei analyzed for each blastomere and condition. **(B)** Single-nucleolus measurements in EMS and P2 nuclei from *sun-1(RNAi)*, *cec-4(ok3124)*; *sun-1(RNAi)*, *zyg-12(RNAi)*, *cec-4(ok3124)*; *zyg-12(RNAi)*, *lmn-1(RNAi)*, and *cec-4(ok3124)*; *lmn-1(RNAi)* embryos. Panels show nucleolar volume and absolute shortest distance to the nuclear envelope measured in 3D. Each point represents one embryo/file mean, calculated from all nucleoli measured within that embryo for the indicated lineage. Boxes indicate the median and middle 50% of embryo/file means, with whiskers showing the full range. Statistical comparisons between matched single and double perturbations were performed on per-embryo means; ns, not significant.

**Figure S3.**
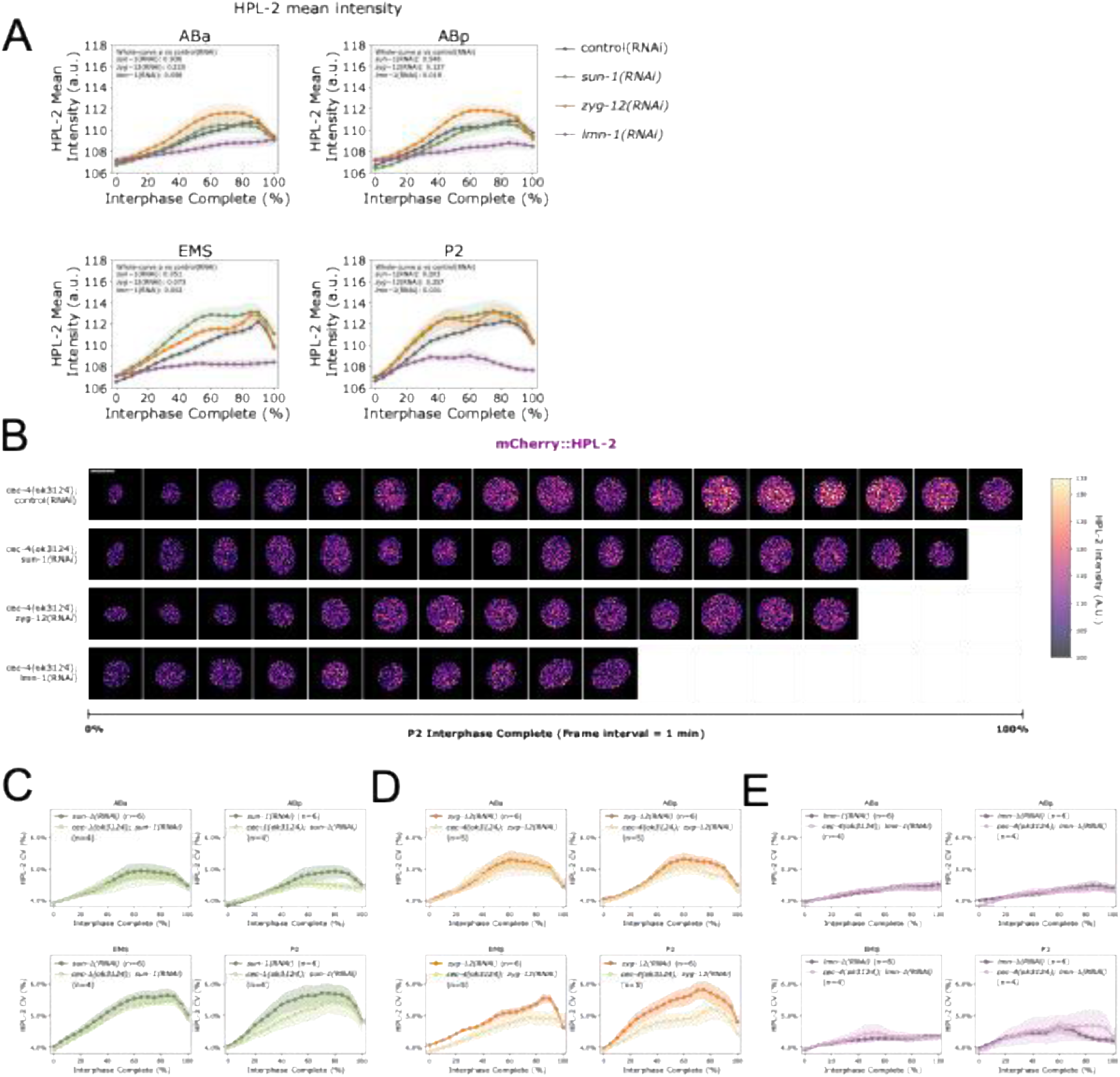
HPL-2 intensity levels and spatial heterogeneity during interphase in single- and double-perturbed embryos. **(A)** HPL-2 mean fluorescence intensity across normalized interphase in ABa, ABp, EMS, and P2 nuclei in control(RNAi), *sun-1(RNAi)*, *zyg-12(RNAi)*, and *lmn-1(RNAi)* embryos. Lines show mean; shading indicates SEM. p values (two-sided Welch’s t-test on whole-curve AUC/100) compare each condition with control(RNAi). **(B)** Representative time series of mCherry::HPL-2 in P2 nuclei during normalized interphase in cec-4(ok3124) embryos with the indicated RNAi conditions. At each displayed time point, the single z-plane containing the maximum FIB-1 signal within the tracked P2 mask was selected independently, and the corresponding mCherry::HPL-2 signal from that same plane is shown. Frame interval, 1 min. Scale bar, 5 µm. **(C-E)** HPL-2 CV in 3D across normalized interphase in ABa, ABp, EMS, and P2 nuclei for each double-perturbed condition compared with its corresponding single RNAi. Lines indicate means; shading indicates SEM. p values: two-sided Welch’s t-test on whole-curve AUC/100.

**Figure S4.**
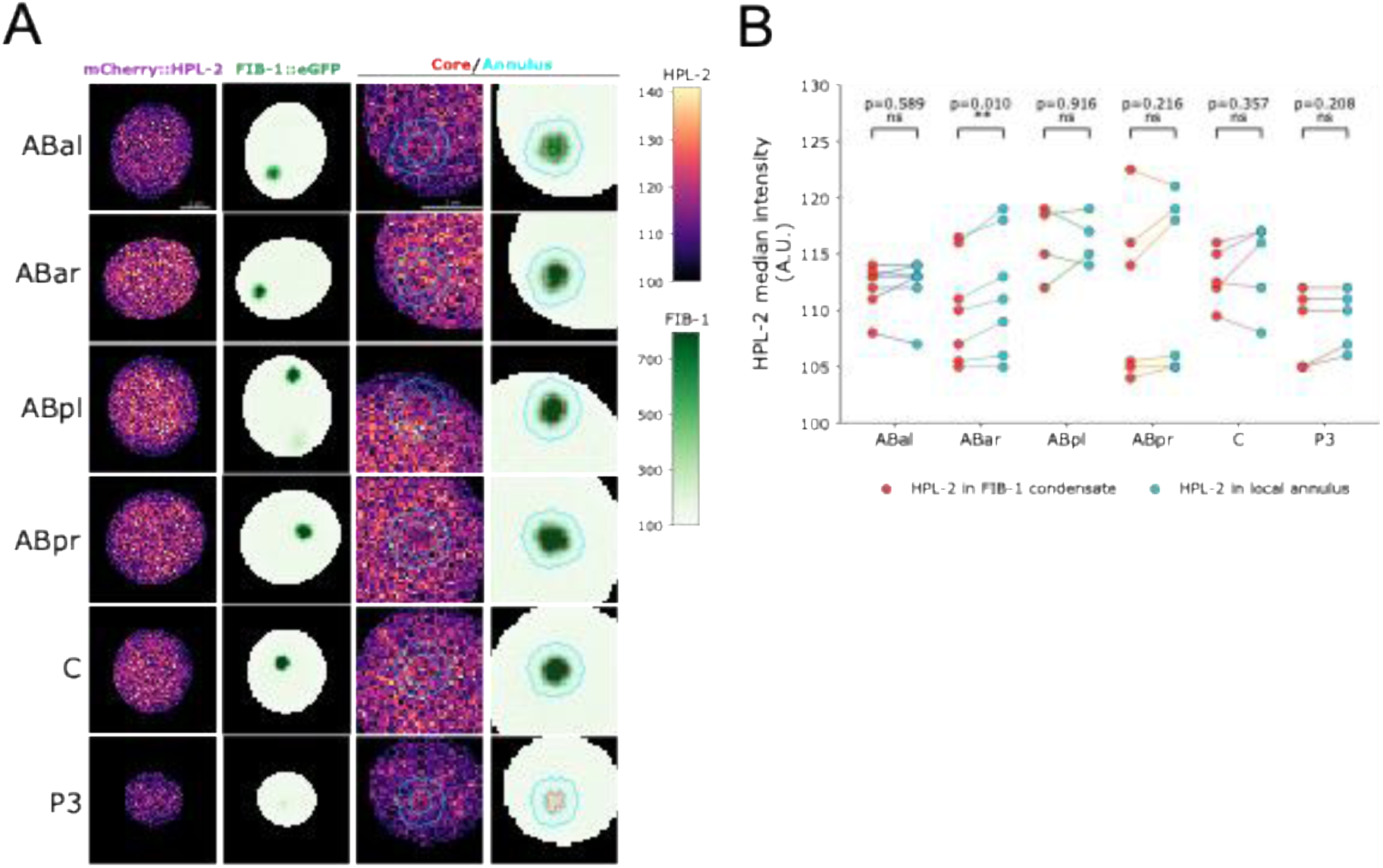
Local HPL-2 signal at FIB-1 condensates in additional 8-cell-stage lineages. **(A**) Representative single-plane mCherry::HPL-2 and FIB-1::eGFP images of wild-type ABal, ABar, ABpl, ABpr, C, and P3 nuclei at the peak FIB-1 frame. Left to right: full HPL-2 plane, corresponding FIB-1 plane, magnified HPL-2 image, and magnified FIB-1 image. Red contours indicate FIB-1 condensate cores; cyan contours indicate the corresponding local annuli. Scale bars, 2 µm. **(B)** Paired median HPL-2 intensity within each FIB-1 condensate core (red points) and its local annulus (cyan points). One condensate per embryo was selected based on the strongest FIB-1 core-to-annulus enrichment; HPL-2 signal was not used for selection. Connected points indicate matched regions from the same nucleus. Exact unadjusted p values were calculated using two-sided paired t-tests and are shown above each lineage; **p < 0.01; ns, not significant. n = 7, 7, 4, 6, 5, and 5 independent embryos for ABal, ABar, ABpl, ABpr, C, and P3, respectively. Corresponding analyses of E and MS are shown in Figs. 5C-D.

**Movie S1. Depleting LINC complexes SUN-1 or ZYG-12 advances nucleolar assembly to 4-cell stage.** Representative time-lapse confocal maximum-intensity projections of FIB-1::eGFP in embryos treated with control(RNAi), *sun-1(RNAi)*, or *zyg-12(RNAi)*. Movies are shown side by side with matched display settings. Time is shown in minutes. The movie pauses at 15 min to highlight precocious, distinct FIB-1 condensates in the EMS and P2 blastomeres at the 4-cell stage in *sun-1(RNAi)* and *zyg-12(RNAi)* embryos, compared with the absence of distinct nucleolar assembly in control(RNAi). Scale bar, 10 µm.

**Movie S2. LMN-1 depletion promotes persistent FIB-1 condensate growth in EMS and P2.** Representative time-lapse confocal maximum-intensity projections of FIB-1::eGFP in embryos treated with control(RNAi) or *lmn-1(RNAi)*. Movies are shown side by side with matched display settings. Time is shown in minutes. The movie pauses at 15 min in control(RNAi) and at 17 min in *lmn-1(RNAi)* to indicate the EMS and P2 blastomeres. At this stage, transient, indistinct FIB-1 condensates in control(RNAi) have already dissolved, whereas FIB-1 condensates in *lmn-1(RNAi)* continue to grow into distinct condensates and persist. Scale bar, 10 µm.

